# Satellite cell expansion is mediated by P-eIF2α dependent *Tacc3* translation

**DOI:** 10.1101/2020.05.13.093302

**Authors:** Ryo Fujita, Graham Lean, Solène Jamet, Steven Hébert, Claudia L. Kleinman, Colin Crist

## Abstract

Translational control of gene expression is an important regulator of adult stem cell quiescence, activation and self-renewal. In skeletal muscle, quiescent satellite cells maintain low levels of protein synthesis, mediated in part through the phosphorylation of eIF2α (P-eIF2α). Pharmacological inhibition of the eIF2α phosphatase with the small molecule sal003 maintains P-eIF2α and permits the expansion of satellite cells *ex vivo*. Paradoxically, P-eIF2α also increases the translation of specific mRNAs, which is mediated by P-eIF2α dependent readthrough of inhibitory upstream open reading frames (uORFs). Here, we ask whether P-eIF2α dependent mRNA translation enables expansion of satellite cells. Using transcriptomic and proteomic analyses, we show a number of genes associated with the assembly of the spindle pole to be upregulated at the level of protein, without corresponding change in mRNA levels, in satellite cells expanded in the presence of sal003. We show that uORFs in the 5’UTR of mRNA for the mitotic spindle stability gene *Tacc3* direct P-eIF2α dependent translation. Satellite cells deficient for TACC3 exhibit defects in expansion, self-renewal and regeneration of skeletal muscle.

**Significance:** Translational control of gene expression has emerged as an important regulator of adult stem cell populations, which maintain low levels of protein synthesis. In adult muscle stem cells, or satellite cells, a portrait of translational control has emerged whereby multiple repression mechanisms prevent the translation of specific mRNAs. It remains unclear how other mRNAs escape repression and are efficiently translated. We show that within the context of low global rates of protein synthesis, satellite cell expansion occurs through the selective translation of *Tacc3* mRNA. *Tacc3* deficient satellite cells expand poorly, leading to defects in skeletal muscle regeneration. Our study provides a more complete picture of translational control of gene expression in adult stem cell populations.

## Introduction

Skeletal muscle regeneration relies on a population of resident adult stem cells, named ‘satellite cells’ for their satellite position around the outside of myofibres and underneath the basal lamina(1). Satellite cells express members of the paired homeodomain family of transcription factors PAX7, and in a subset of muscle, PAX3(2). Satellite cells are mitotically quiescent and activate the myogenic program and the cell cycle in response to muscle injury. Activated satellite cells exhibit remarkable proliferative capacity to rapidly expand the population of myogenic progenitors required to efficiently regenerate muscle. Moreover, this proliferative phase is marked by symmetric cell divisions which drive expansion of the satellite cell pool and asymmetric cell divisions, mediated in part through an EGFR-AuroraA kinase signaling pathway, which ensure satellite cell self-renewal (3). How satellite cells achieve this proliferative capacity, while maintaining the fidelity of cell division, remains unclear.

Quiescent satellite cells have few activated mitochondria and generate low levels of ATP(4, 5). To manage available energy resources, quiescent satellite cells likely maintain low rates of protein synthesis, mediated in part by the phosphorylation of translation initiation factor eIF2α (eIF2α)(6). Genetic inactivation of eIF2α phosphorylation (P-eIF2α) in satellite cells leads to increased global protein synthesis, activation of the myogenic program, and failure to self-renew. Pharmacological inhibition of eIF2α dephosphorylation with the small molecule sal003(7, 8)maintains low levels of protein synthesis and enables *ex vivo* expansion of cultured satellite cells that retain their regenerative potential(6, 9).

While maintaining low levels of protein synthesis, quiescent satellite cells also repress the translation of specific transcripts that maintain satellite cells ‘primed’ to activate the myogenic program and the cell cycle. *Myf5* transcripts are repressed by miR-31 and accumulate in cytoplasmic RNA granules(10), which require P-eIF2α for their assembly and maintenance(6). *MyoD* transcripts are repressed by the activity of RNA binding proteins TTP(11) and STAUFEN1(12), and transcripts required to activate the cell cycle, such as *Dek*, are repressed by miR-489(13).

While the role of post-transcriptional mechanisms repressing gene expression in satellite cells is well documented, a more complete picture requires that we also understand how specific mRNAs escape repression and are translated efficiently. When cells are under stress, P-eIF2α dependent mRNA translation is illustrated by selective translation of mRNAs for *Atf4* and *Chop*, which are required to initiate an integrated stress response (ISR) (14, 15). These P-eIF2α dependent mRNAs have inhibitory upstream open reading frames (uORFs) that prevent the translation of the main ORF under normal conditions, while stress-induced P-eIF2α permits the readthrough of the uORFs to enhance the translation of the main ORF required for the ISR.

We propose a model by which P-eIF2α dependent mRNA translation regulates satellite cell quiescence, self-renewal and expansion. In this study, we determined how satellite cells modify their transcriptome and proteome while expanded with sal003. We show that sal003 pushes satellite cell gene expression towards a progenitor cell phenotype, while preventing differentiation. By focusing on genes upregulated at the level of protein, without a corresponding increase in mRNA, we demonstrate that satellite cell expansion is mediated in part through P-eIF2α dependent translation of mRNA for *Tacc3* (transforming acidic coiled coil protein 3), an AuroraA kinase substrate that is essential for microtubule assembly, maintenance of the spindle pole and ensuring the fidelity of cell division (16, 17).

## Results

### Pharmacological inhibition of eIF2α dephosphorylation leads to expansion of satellite cells with distinct transcriptional profiles

To determine the mechanism of action of sal003, and simultaneously ask how P-eIF2α enables satellite cell expansion, we used RNA-seq to determine global changes in gene expression in satellite cells isolated from diaphragm and abdominal muscle of adult *Pax3^GFP/+^* mice after four-day culture in the presence of 10μM sal003 (Supplementary Data 1). Principal component analysis (PCA) reveals a strong, global effect on the transcriptome, with sal003 treatment explaining 90% of the variance in the data (Fig. 1A). Consistent with the effect of sal003 to reduce global protein synthesis, we observed a general trend towards downregulation of genes that produced an asymmetrical volcano plot (Fig. 1B), as well as larger number of downregulated genes (958 vs 383, p-value < 0.00001). We generated a z-score heatmap to visualize selected genes that are upregulated or downregulated at least 2-fold in the presence of 10μM sal003 (Fig. 1C).

**Fig. 1.**
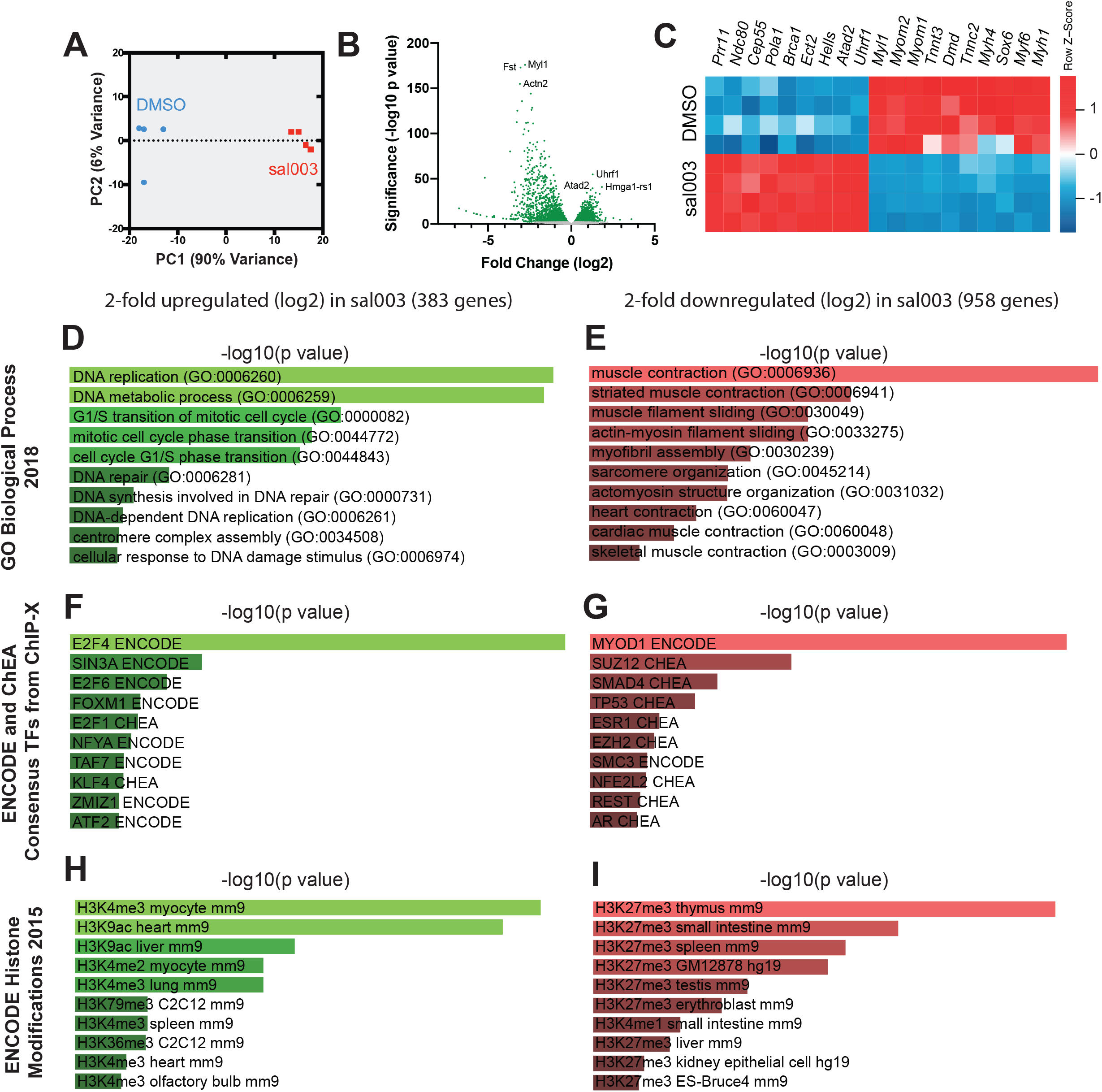
Transcriptome analysis of satellite cells cultured in the presence of 10μM sal003. (*A*) A principal component analysis (PCA) plot derived from RNA-seq transcriptome profiling of satellite cells cultured under normal conditions (DMSO, blue) and 10μM sal003 (sal003, red). (*B*) A volcano plot depicting differentially expressed genes (mRNA) between satellite cells cultured in DMSO (control) and 10μM sal003. The threshold of p value 0.05 is indicated by green and grey data points. The top three significantly downregulated and upregulated genes are indicated. (*C*) A z-score heatmap of selected differentially expressed genes in satellite cells after 4-day culture in 10μM sal003. (*D-I*) Analysis of upregulated (*D, F, H*) and downregulated (*E, G, I*) genes were performed by gene set enrichment analysis for GO Biological Process (*D, E*), ENCODE and ChEA Consensus Transcription Factors (*F, G*) and ENCODE Histone Modifications (*H, I*). Intensity and length of bars indicate significance (−log10 p value). The top 10 enriched gene sets are shown in each category.

To understand the mechanisms underlying changes in gene expression when satellite cells are cultured in the presence of sal003, we used Enrichr (18) to perform gene ontology (GO) analyses (GO Biological Process 2018), as well as to identify enrichment of consensus target genes for transcription factors (ENCODE and ChEA Consensus TFs from ChIP-X) and histone modifications (ENCODE Histone Modifications 2015). Amongst the 383 genes upregulated greater than 2-fold (log2>1), there is strong enrichment for GO Biological Process gene sets related to DNA replication, mitosis and cell cycle regulation (Fig. 1D). When satellite cells are cultured in the presence of sal003, we observe enrichment for target genes of known transcriptional regulators of myogenic progenitor survival and proliferation (E2F1, SIN3, E2F6, E2F4, FOXM1) (19) (Fig. 1E). Upregulated genes also show enrichment for chromatin marks associated with actively transcribed genes in myogenic progenitors (H3K4me3, H3K9ac, H3K6me3) (Fig. 1F).

Amongst the 958 genes downregulated greater than 2-fold (log2< −1) there is strong enrichment for GO Biological Process gene sets related to differentiated skeletal muscle, including muscle contraction, muscle filament sliding, myofibril assembly and sarcomere organization (Fig. 1G). Amongst these genes, consensus target genes for transcription factor MYOD is predominant (Fig. 1H) and moreover, down-regulated genes are enriched for chromatin marks common to repressed gene bodies (H3K27me3) in non-skeletal muscle tissue (Fig. 1I).

### Pharmacological inhibition of eIF2α dephosphorylation modifies the satellite cell proteome

Since sal003 inhibition of P-eIF2α dephosphorylation is expected to impact rates of mRNA translation independent of changes in transcription, we simultaneously asked how sal003 modifies the satellite cell proteome. We cultured satellite cell under normal conditions or in the presence of sal003, then labeled cell lysates with tandem mass tag (TMT) reagents to identify and quantify changes in protein expression by high resolution mass spectrometry (Fig. 2A, Supplementary Data 2).

**Fig. 2.**
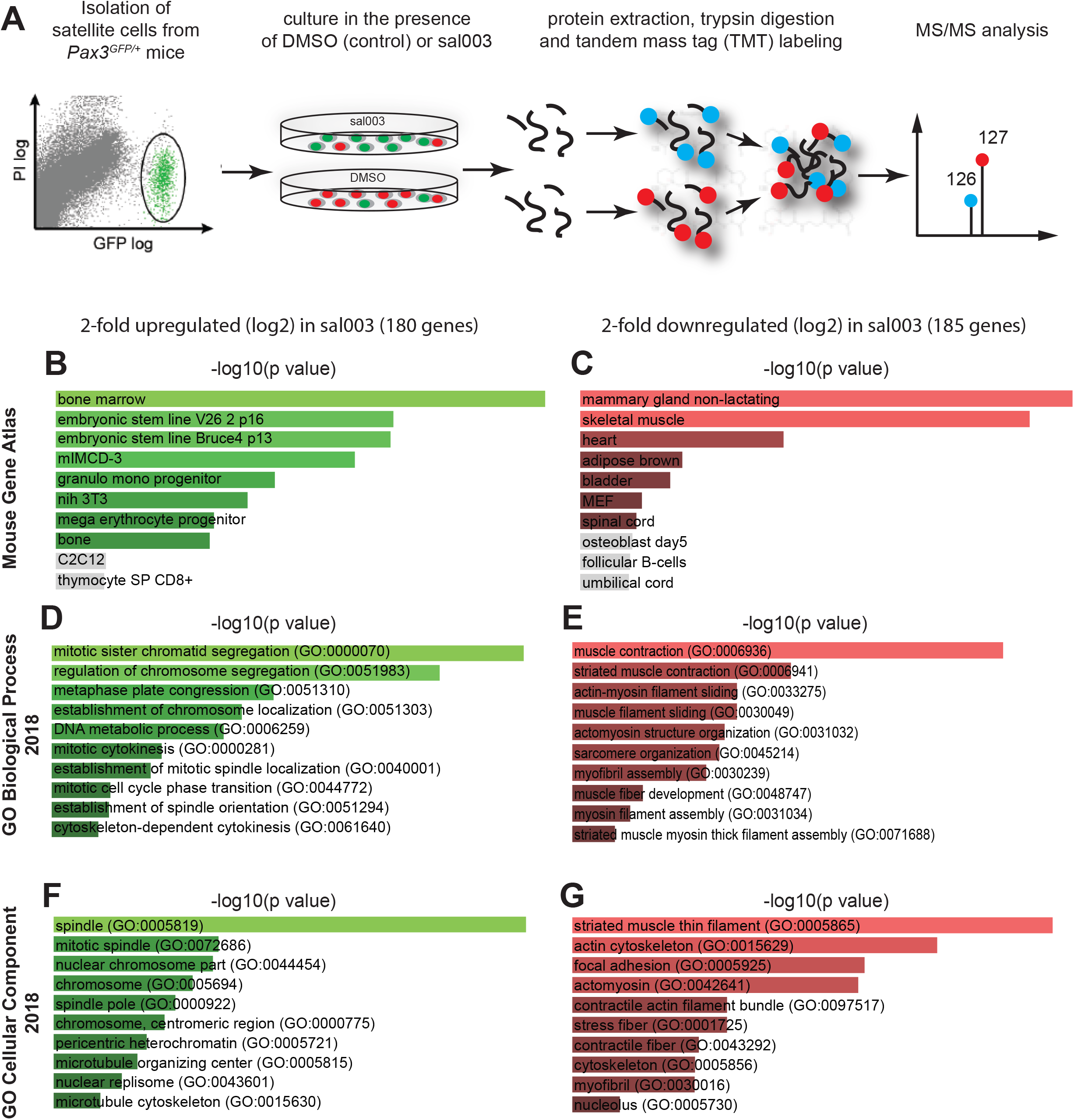
Proteome analysis of satellite cells cultured in the presence of 10μM sal003. (*A*) Schematic representation of satellite cell proteome analysis by TMT labeling and mass spectrometry. (*B-G*) Analysis of upregulated (*B, D, F*) and downregulated (*C, E, G*) genes performed by gene enrichment analysis for tissue expression in the Mouse Gene Atlas (*B, C*), GO Biological Process (*D, E*) and GO Cellular Component (*F, G*). Intensity and length of bars indicate significance (−log10 p value). Grey bars, p<0.05. The top 10 enriched gene sets are shown in each category.

Of the 180 genes upregulated 2-fold or greater (log2>1) quantified by mass spectrometry, there is strong enrichment for genes expressed in mouse progenitor and stem cell lines (BioGPS, Fig. 2B). Upregulated genes are enriched in chromatid and chromosome separation (GO Biological process, Fig. 2D), and are associated with the mitotic spindle, centromere and microtubules (GO Cellular Component, Fig. 2F). In contrast, the presence of 10μM sal003 in satellite cell culture conditions down-regulates 185 genes (2-fold, log2<−1) quantified by mass spectrometry. These genes are enriched in mouse skeletal muscle tissue (BioGPS, Fig. 2C), are associated with muscle contraction, muscle filament sliding (GO Biological Processes, Fig. 2E) and are components of striated muscle thin filament, actin cytoskeleton, focal adhesions and actomyosin (GO Cellular Component, Fig. 2G).

To reveal transcripts potentially translated in a P-eIF2α dependent manner, we identify 140 genes that are upregulated at the level of protein (mass spectrometry, log2>1), without a corresponding increase reported by mRNA (RNA-seq, log 2<1) (Fig. 3A, B). Upregulated genes include inhibitors of myogenic differentiation (PAX7, ID3, CABIN1) and chromatin modifiers (KDM5C, KMT2A, SETD1A, SETD2) that may account for, in part, transcriptional changes in gene expression observed (Fig. 1). The most significantly represented class of genes are those involved in spindle assembly (TACC3, CDC20, TPX2, NEDD1, RACGAP1, ESPL1, SPAG5, INCENP, VPS4B, KIF2C, SKA3) (Fig. 3A, B, Supplementary Data 3).

**Fig. 3.**
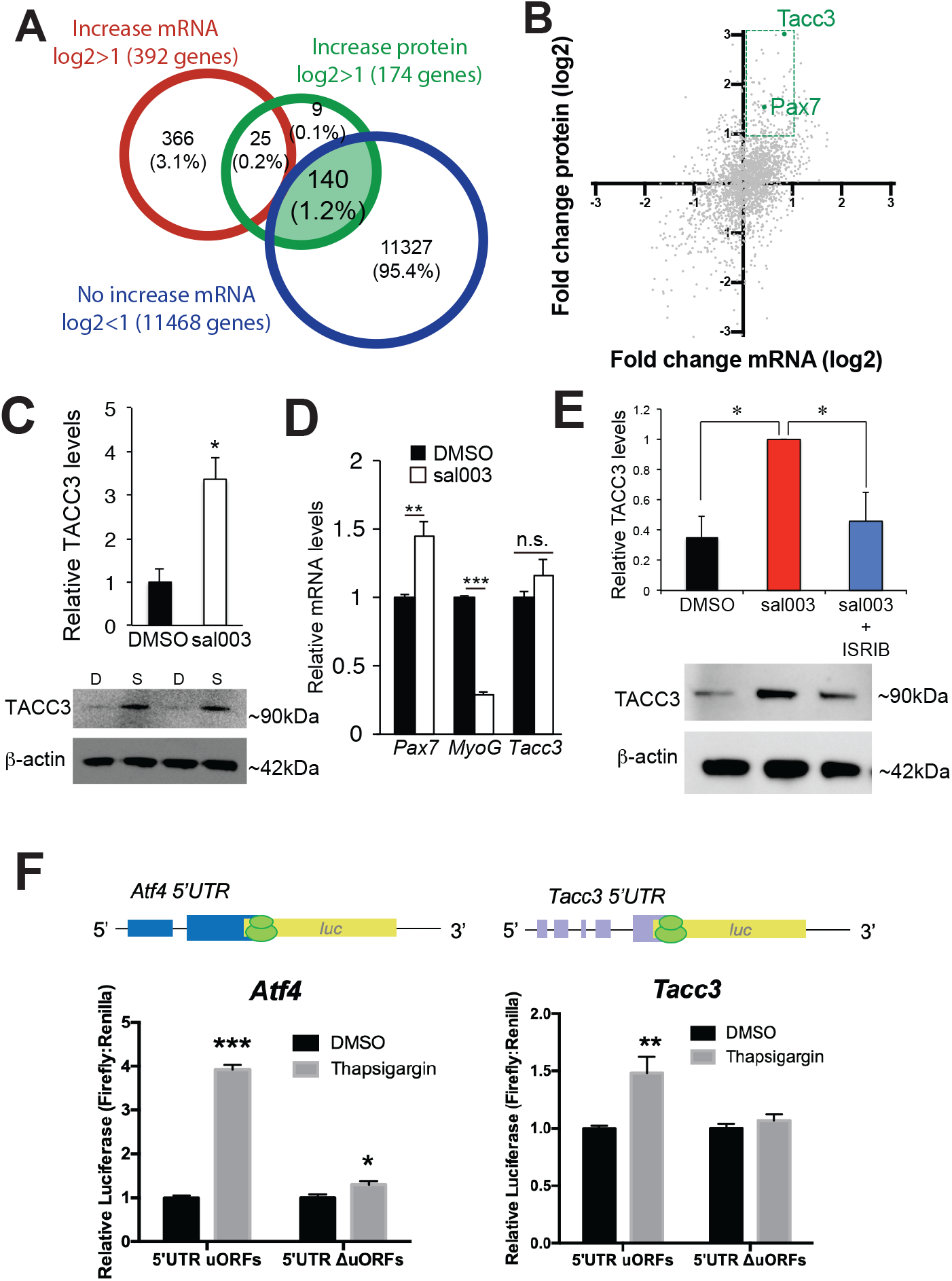
Satellite cells upregulate protein levels independent of mRNA when cultured in sal003: P-eIF2α dependent Tacc3 mRNA translation. (*A*) Venn diagram showing the relationship of genes upregulated at the level of mRNA (red) and protein (green) in satellite cells cultured in 10μM sal003. 140 genes (green shading) are upregulated at the level of protein (mass spectrometry, log2>1), without a corresponding increase reported by mRNA (RNA-seq, log 2<1) (blue). (*B*) Scatterplot of gene expression reported by transcriptome (x-axis) and proteome (y-axis) analyses. PAX7 and TACC3 are highlighted (green) within the subset of genes upregulated at the level of protein (mass spectrometry, log2>1), without a corresponding increase reported by mRNA (RNA-seq, log 2<1) (green dotted line). (*C*) Immunoblotting against TACC3 and β-actin from cell lysates of *Pax3^GFP/+^* satellite cells cultured in the presence of DMSO (D, control) and sal003 (S). Relative levels of TACC3 normalized to b-actin are indicated, with representative immunoblots shown. (*D*) Relative *Pax7*, *MyoG* and *Tacc3* levels, determined by RT-qPCR, after four day culture in the presence of DMSO (control) or 10μM sal003. Levels are reported normalized to *actb* and relative to DMSO conditions. *(E)* Immunoblotting against TACC3 and β-actin from cell lysates of *Pax3^GFP/+^* satellite cells cultured in the presence of DMSO, sal003 or sal003 with ISRIB (200nM). Relative levels of TACC3 normalized to β-actin are indicated, with representative immunoblots shown. (*F*) Relative Firefly:Renilla luciferase activity in 293 cells transfected with pGL3-Atf4-luc, pGL3-Atf4mut-luc, pGL3-Tacc3-luc or pGL3-Tacc3mut-luc (firefly), and pRL-TK (renilla) after overnight culture in the presence of 1 μM thapsigargin (TG). Values indicate mean (n = 3) ± SEM. * p < 0.05, ** p < 0.01, *** p < 0.001.

### P-eIF2α dependent translation of *Tacc3* mRNA

We further focused our attention on Transforming acidic coiled coil protein 3 (TACC3), which was the most significantly upregulated gene reported by mass spectrometry and is a representative gene of the most significantly enriched gene ontology (spindle pole). *Tacc3* mRNA contains 5 uORFs in its 5’UTR, potentially enabling selective translation of the main ORF for *Tacc3* by P-eIF2α. TACC3 is a substrate of AuroraA kinase and functions at the centrosome to regulate microtubule nucleation, promote stability of the spindle apparatus and the fidelity of cell division(16, 17). Inhibition of TACC3 by knockout, knockdown and pharmacological strategies reveal a role for TACC3 in maintaining or expanding adult and cancer stem cell populations, while it is dispensable for stem cell differentiation(20–23).

First, we confirm that 4-day culture of satellite cells in the presence of 10μM sal003 increases TACC3 protein levels (Fig. 3C), without a corresponding change in *Tacc3* mRNA (Fig. 3D), validating the identification of *Tacc3* in our global gene expression profiles (Fig. 3A, B). Next, we cultured satellite cells in the presence of sal003 to induce P-eIF2α levels and in the presence of ISRIB, which bypass the effect of P-eIF2α by promoting eIF2-GTP recycling independent of eIF2α(24). The additional treatment of satellite cells with ISRIB lowers TACC3 protein levels to normal culture conditions, further linking *Tacc3* mRNA to P-eIF2α dependent translation (Fig. 3E).

To ask whether uORFs present in the 5’UTR of *Tacc3* mRNA mediate P-eIF2α dependent translation in a manner similar to *Atf4*, we cloned the 5’UTRs of these two mRNAs upstream of a luciferase reporter and generated additional reporters whereby upstream ORFs are eliminated by mutation of the ATG start codon. We transfected 293 cells with *Atf4* and *Tacc3* P-eIF2α reporters, and further cultured 293 cells under normal conditions or in the presence of thapsigargin (TG) to induce high P-eIF2α levels(15). P-eIF2α dependent translation of the luciferase reporter occurs in the presence of TG when uORFs in *Atf4* or *Tacc3* are present, but not when these uORFs are eliminated by mutation of the start codon (Fig. 3F).

### Tacc3 is required for expansion of self-renewing satellite cells *ex vivo*

Since sal003 upregulates TACC3 and facilitates *ex vivo* expansion of self-renewing satellite cells, we next asked whether *Tacc3* is required for satellite cell self-renewal. We examined satellite cells from *Pax7^CreERT2/+^*; *Tacc3^fl/fl^* mice(22, 25, 26), such that tamoxifen (tmx) treatment would result in loss of TACC3 from PAX7-expressing satellite cells. First, satellite cells were isolated from tamoxifen treated *Pax7^CreERT2/+^*; *Tacc3^fl/fl^* mice and further cultured in the presence of 4-Hydroxytamoxifen (4-OHT) to eliminate *Tacc3* expression in cultured satellite cells (Fig. 4A-C). In the absence of TACC3, PAX7 is present diffusely within the nucleus and cytoplasm (Fig. 4B, D), and low PAX7 levels are confirmed by western blotting (Fig. 4E) and RT-qPCR (Fig. 4F). Satellite cells defective for *Tacc3* have a pronounced defect to expand into large myocolonies *ex vivo* (Fig. 4G, H). Further analysis of myocolonies present after 4 day culture indicate precocious differentiation, as revealed by an increase in PAX7-negative (−), MYOD-positive (+) nuclei, and a decrease in self-renewal, as indicated by the near absence of PAX7(+), MYOD(−) nuclei representing the ‘reserve cell’ population (Fig. 4G, I). Next, we rescued TACC3 expression in satellite cells isolated from tmx treated *Pax7^CreERT2/+^*; *Tacc3^fl/fl^* mice with lentiviral vectors driving *Tacc3* expression under the phosphoglycerate kinase 1 (PGK) promoter (Fig. 4J, K). Increased TACC3 levels led to more robust PAX7 expression and localization of PAX7 to the nucleus (Fig. 4K, L).

**Fig. 4.**
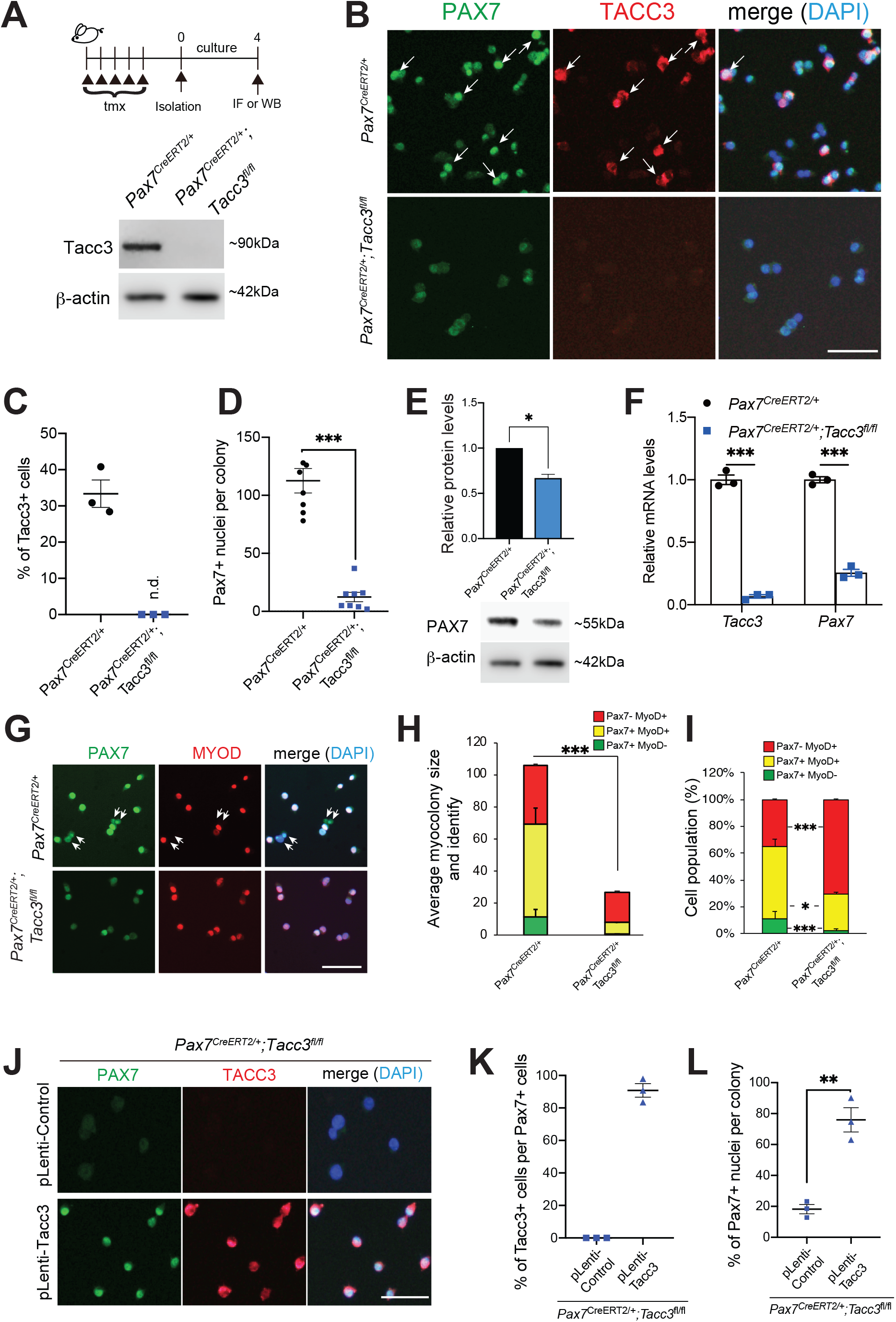
TACC3 is required for satellite cell expansion and self-renewal *ex vivo*. (*A-B*) Isolation of satellite cells from tamoxifen (tmx) treated *Pax7^CreERT2/+^*; *Tacc3^fl/fl^* mice, followed by 4 day culture in the presence of 4-hydroxytamoxifen (4-OHT) leads to the absence of TACC3 reported by (*A*) immunoblotting of lysates with antibodies against TACC3 and β-actin and (*B*) immunofluorescence of satellite cell cultures with antibodies against PAX7 (green) and TACC3 (red). Arrows (white) indicate presence of TACC3(+) satellite cells that are strongly immunolabeled for PAX7. (*C*) Quantification of TACC3 positive nuclei in (B), after 4-day culture of satellite cells isolated from tmx treated *Pax7^CreERT2/+^* and *Pax7^CreERT2/+^*; *Tacc3^fl/fl^* mice in (B). (*D*) Numbers of PAX7(+) nuclei in (B). (*E*) Immunoblotting of satellite cell lysates with antibodies against PAX7 and β-actin after 4-day culture of satellite cells isolated from tmx treated *Pax7^CreERT2/+^* and *Pax7^CreERT2/+^*; *Tacc3^fl/fl^* mice. (*F*) RT-qPCR analysis of *Tacc3* and *Pax7* expression after 4 day culture of satellite cells isolated from tmx treated *Pax7^CreERT2/+^* and *Pax7^CreERT2/+^*; *Tacc3^fl/fl^* mice. (*G*) Immunofluorescence analysis with antibodies against PAX7 (green) and MYOD (red) after 4 day culture of satellite cells isolated from tmx treated *Pax7^CreERT2/+^* and *Pax7^CreERT2/+^*; *Tacc3^fl/fl^* mice. (*H*) Quantification of myocolony size in (G). (*I*) Quantification of satellite cells undergoing self-renewal (reserve cell population, PAX7(+), MYOD(−)), activation (PAX7(+), MYOD(+)) and differentiation (PAX7(−), MYOD(+) in (G). (*J*) Immunofluorescence analysis with antibodies against PAX7 (green) and TACC3 (red) after 4-day culture of satellite cells isolated from tmx treated *Pax7^CreERT2/+^*; *Tacc3^fl/fl^* mice and transduced with lentiviral vectors overexpressing *Tacc3 (pLenti-Tacc3)*. (*K*) Numbers of PAX7(+), TACC3(+) cells in (J). (*L*) Numbers of PAX7(+) nuclei in (J). Scale bars represent 50μm (B, G and J). Values indicate mean (n = 3) ± SEM. * p < 0.05, ** p < 0.01, *** p < 0.001.

The importance of TACC3 to promote satellite cell expansion, rates of proliferation, and satellite cell self-renewal, while preventing precocious differentiation, is also confirmed by siRNA knockdown of *Tacc3* (Supplementary Fig. 1) and by TACC3 inhibition with the small compound KHS101(21) (Supplementary Fig. 2).

### *Tacc3* deficient satellite cells expand poorly, leading to defects in muscle regeneration

TACC3 is expressed in a subset of quiescent satellite cells *in vivo*, where it is present diffusely in the cytoplasm (Fig. 5A). In activated satellite cells that remain associated with single EDL myofibres, TACC3 appears to localize to perinuclear areas and to spindle poles (Fig. 5B). To reveal a role for TACC3 in satellite cells *in vivo*, we inactivated *Tacc3* by tmx administration to *Pax7^CreERT2/+^*; *Tacc3^fl/fl^* mice. After five days of tmx, numbers of PAX7(+) nuclei along single isolated EDL myofibres remain unchanged (Fig. 5C). Given TACC3’s role in mitotic spindle assembly and maintaining the fidelity of cell division(16, 17), we asked whether TACC3 was required for expansion of activated satellite cells *in vivo*. We used cardiotoxin (ctx) to injure *tibialis anterior* (TA) muscle of tmx treated *Pax7^CreERT2/+^*; *Tacc3^fl/fl^* mice (Fig. 5D). Seven days after acute injury, we confirm decreased numbers of TACC3(+) and PAX7(+) satellite cells (Fig. 5D-F) and the decrease in PAX7 expression is confirmed by western blotting of lysates of isolated satellite cells (Fig. 5G). The inability of *Tacc3*-deficient satellite cells to expand *in vivo* should lead to defects in muscle regeneration, which is illustrated by decreased myofiber size 21 days after ctx injury (Fig. 5H-I).

**Fig. 5.**
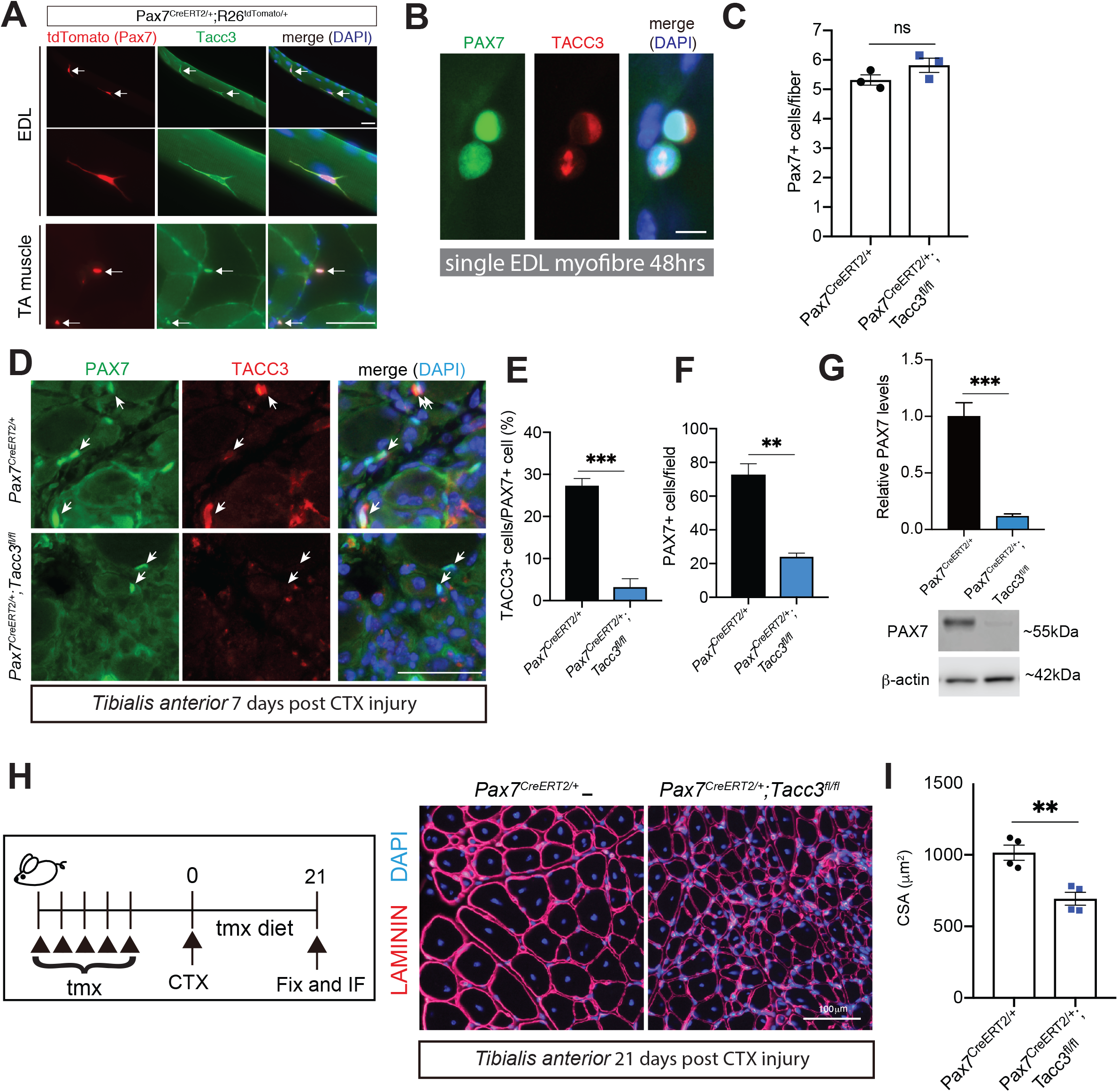
TACC3 is required for satellite cell expansion and skeletal muscle regeneration *in vivo*. (*A*) Tdtomato fluorescence (PAX7, red) combined with immunofluorescence analysis with antibodies against TACC3 (green) on single EDL myofibres and transverse TA sections from tamoxifen treated *Pax7^CreERT2/+^*; *R26^tdTomato/+^* mice. (*B*) Immunofluorescence analysis with antibodies against PAX7 (green) and TACC3 (red) on single EDL myofibres after 48-hour culture. (*C*) Numbers of PAX7(+) satellite cells immediately after tmx treated *Pax7^CreERT2/+^* and *Pax7^CreERT2/+^*; *Tacc3^fl/fl^* mice. (*D*) Immunofluorescence analysis with antibodies against PAX7 (green) and TACC3 (red) on transverse sections of TA muscle from tmx treated *Pax7^CreERT2/+^* and *Pax7^CreERT2/+^*; *Tacc3^fl/fl^* mice, 7 days after ctx injury. Arrows (white) indicate the position of PAX7(+) nuclei. (*E*) Quantification of numbers of TACC3(+), PAX7(+) satellite cells in (D). (*F*) Quantification of PAX7(+) nuclei per field of view in (D). (*G*) Immunoblotting of lysates obtained from MACS sorted satellite cells, 7 days after ctx injury, with antibodies against PAX7 and β-actin. Quantification of PAX7 levels, relative to β-actin levels by densitometry of n=3 independent experiments, with a representative immunoblot shown. (*H*) Immunofluorescence analysis with antibodies against LAMININ (red) counterstained with DAPI (blue) on transverse sections of TA muscle from tmx treated *Pax7^CreERT2/+^* and *Pax7^CreERT2/+^*; *Tacc3^fl/fl^* mice, 21 days after ctx injury. (*I*) Quantification of myofiber cross section area (CSA) in (H). Scale bars represent 50μm (A, D), 10μm (B) or 100μm (H). Values indicate mean (n = 3) ± SEM, except in (I) where (n=4). ** p < 0.01, *** p < 0.001, ns, not significant.

## Discussion

### P-eIF2α dependent *Tacc3* translation permits satellite cell expansion *ex vivo*

In this work, we set out to understand how sal003-mediated inhibition of eIF2α dephosphorylation enables the expansion of satellite cells and further identify important regulators of satellite cell behaviour during a phase of expansion. Phosphorylation of eIF2α is required for satellite cell quiescence since genetic tools to eliminate eIF2α phosphorylation cause satellite cells to activate, enter the cell cycle and contribute to differentiation, while being defective for self-renewal. The presence of sal003 in culture conditions leads to decreased rates of proliferation, observations that are consistent with a role for P-eIF2α in quiescent satellite cells(6). We propose that expansion occurs because slowly proliferating satellite cells avoid differentiating into myoblasts that become post-mitotic and fuse to form the myofiber. Our data supports this model because genes required for differentiated skeletal muscle tissue are the most significantly downregulated in the presence of 10μM sal003 (Fig. 1, 2).

Here, we asked whether sal003 enables satellite cell expansion by P-eIF2α dependent translation of specific mRNAs. Culture of satellite cells in the presence of sal003 broadly leads to maintenance of gene expression associated with stem and progenitor cells (Fig. 1, Fig. 2). Amongst our candidates for P-eIF2α dependent translation, enriched genes were most commonly associated with the spindle pole, centrosome and microtubule assembly (Fig. 3, Supplementary Table 1), suggesting that sal003 permits the translation of mRNAs required to maintain the fidelity of cell division. Amongst these genes, we focused on *Tacc3* because its 5’UTR includes 5 uORFs and we demonstrate readthrough of these inhibitory uORFs in a P-eIF2α dependent manner. Moreover, *Tacc3* depletion leads to a loss of PAX7 expression in satellite cells and precocious differentiation, while *Tacc3* overexpression leads to increased expression of PAX7 in satellite cells. Whether PAX7-expression is directly or indirectly affected by TACC3 remains unclear, but we note that TACC3 regulates subcellular localization and physical association of transcription factors to control hematopoeiesis(27)

### Satellite Cell Expansion and Self-Renewal *Ex Vivo* and *In Vivo* Requires TACC3

TACC3 is amongst several substrates of AuroraA kinase that influence mitotic spindle assembly(17). An EGFR-AuroraA kinase signaling pathway orients the mitotic spindle apicobasally to facilitate asymmetric cell divisions to maintain the satellite cell pool(3). It is unknown whether TACC3 regulates the orientation of the mitotic spindle, but we show that TACC3 depletion preferentially reduces the ‘reserve cell’ population of satellite cells undergoing self-renewal *ex vivo*. Failure to expand a population of PAX7(+) progenitors is also observed *in vivo* 7 days after ctx injury, which ultimately leads to a defect in skeletal muscle regeneration at later timepoints (21 days).

### TACC3 Accumulates in Quiescent Satellite Cells

We also demonstrate that TACC3 marks a subset of quiescent satellite cells *in vivo*, where it is present diffusely throughout the cytoplasm. Future studies will ask whether TACC3 directly regulates quiescent satellite cells. Additional roles for TACC3 outside the context of the centrosome or spindle pole have been derived from studies in *Xenopus laevis*, where the *Tacc3* homolog is *Maskin*. In developing *Xenopus* oocytes, Tacc3/Maskin regulates the translation of mRNAs with cytoplasmic polyadenylation elements (CPE), such as *cyclinB* mRNA, to prevent cell cycle progression(28–30). The regulation of *cyclinB* mRNA translation would have clear implications for maintaining quiescence, but to our knowledge TACC3 has not yet been characterized as a translational regulator of CPE containing mRNAs in mammalian cells. TACC3 (Maskin) is also required for the assembly of the microtubule network that underlies axonal outgrowth in developing *Xenopus* neurons(31) and for viral and cellular cargo transport in human cells(32).

Increasing evidence points to translational control of gene expression as an important regulator of adult stem cell quiescence and self-renewal, with multiple mechanisms mediated by microRNA and RBPs repressing the translation of specific mRNAs. Here, we show increases in protein production, independent of mRNA levels, for a number of genes expressed in satellite cells cultured in the presence of sal003. We demonstrate P-eIF2α dependent translation of mRNA for *Tacc3* and show a requirement for *Tacc3* expression in expanding satellite cells *ex vivo* and *in vivo*. Our findings suggest an additional role for P-eIF2α dependent translation of mRNA in the maintenance of adult stem cell populations and outside the context of the ISR.

## Materials and Methods

### Mice

Animal care practices were in accordance with the federal Health of Animals Act, as practiced by McGill University. All mice were maintained on a C57/Bl6 background. *Tacc3^fl/fl^* mice were kindly provided by R. Yao (22). *Pax7^CreERT2/+^* (25) and *Rosa26^tdTomato^* (33)mice were obtained from Jackson Laboratories. Tmx (Cayman Chemical) was administered in corn oil, 30% ethanol by intraperitoneal injections (2.5 mg/day) for 5 days and when indicated, mice were maintained on a tmx diet (80 mg/kg body weight/day, Envigo). For muscle regeneration, 8-week-old mice were anesthetized by isofluorane (CDMV) inhalation and 50 μl of 10 μM ctx (Sigma) was injected into the TA muscle. At indicated timepoints, muscle was harvested for analysis by immunofluorescence.

### Cell and Single-Fiber Isolation and Culture

Satellite cells were isolated from abdominal and diaphragm muscle of 8 week old *Pax3^GFP/+^*, *Pax7^CreERT2/+^*, *R26^tdtomato/+^*; *Pax7^CreERT2/+^*, *R26^tdtomato^*; *Tacc3^fl/fl^* mice by flow cytometry (GFP or tdTomato) as previously described(6) using a FACSAriaII cell sorter (BD Biosciences), or alternatively from *Pax7^CreERT2/+^* or *Pax7^CreERT2/+^*, *Tacc3^fl/fl^* mice by MACS Satellite Cell Isolation Kit, together with anti-Integrin α-7 MicroBeads (Miltenyi). Single myofibres were isolated by trituration of 0.5% collagenase D (Sigma)-treated EDL muscle of 8-week old adult mice. Cells and single EDL myofibres were cultured in 39% DMEM, 39% F12, 20% fetal calf serum (FCS) (Life Technologies), 2% UltroserG (Pall Life Sciences). When indicated, culture conditions were supplemented with 0.1% dimethylsulfoxide (DMSO control, Sigma), 10μM sal003 (Sigma), 200nM ISRIB (Cayman), 10 μM 4-OHT (Cayman) or 2.5μM KHS101 (Sigma). For siRNA experiments, satellite cells were transfected after 48h culture with siRNAs against *Tac*c3 (20nM) or Mission^®^ siRNA Universal Negative control (Sigma) with jetPRIME^®^ transfection reagent (Illkirch-Graffenstaden, France). Transfected satellite cells were cultured for 2 additional days. For 5-ethynyl-2’-deoxyuridine (EdU) incorporation assays (Life Technologies), 10μM EdU was included in culture for two hours. 293 cells were cultured in 90% DMEM, 10% FCS and supplemented when indicated with 1 μM thapsigargin (TG, Sigma).

### Tacc3 Overexpression

The mouse *Tacc3* sequence was amplified by PrimeSTAR^®^ Max DNA polymerase (TAKARA) using forward 5’-CTCCCCAGGGGGATCATGAGTCTGCATGTCTTAAAT-3’ and reverse 5’-GAGGTTGATTGTCGATCAGATCTTCTCCATCTTAG-3’ primers. The purified fragment was cloned into *Bam*HI-*Sal*I site of pLenti-PGK-GFP(34) with In-Fusion cloning HD kit (TAKARA). HEK293T cells were transfected with the pLenti-PGK-Tacc3 or pLKO.1 TRC (control, Addgene 10879), pMD2G (Addgene 12259) and psPax2 (Addgene 12260) with jetPRIME^®^ transfection reagent (Illkirch-Graffenstaden, France). Six hours later, the medium was changed and 42 hours later, supernatant was collected, filtrated through a 0.45μm filter, and concentrated with a Lenti-X™ Concentrator (TAKARA). Titers (~1×10^8^ infectious units/ml) were calculated by GFP analysis of transduced 293T cells. Satellite cells isolated by either MACS or FACS described above (1×10^4^ cells/35mm dish or a well of 6 well plate) were transduced with 4 μl of lentivirus solution with polybrene (5μg/ml). Twenty-four hours after transduction, the lentivirus containing medium was carefully removed and replaced with fresh satellite cell medium. Transduced satellite cells were cultured for 3 additional days.

### Luciferase assay

The *Atf4*- and *Tacc3*-luciferase constructs were made by cloning gBlock gene fragments (Integrated DNA Technologies) corresponding to 5’UTRs of *mus musculus Atf4* and *Tacc3*, fused to the 5’end of the *fluc* gene up to the *Nar*I restriction enzyme site. Mutant versions of *Atf4* and *Tacc3* 5’UTRs were designed with each ATG start codon of uORFs deleted. These 5’UTR gene fragments were cloned into *Hind*III, *Nar*I sites upstream of the firefly luciferase (*fluc*) gene in pGL3-promoter plasmids, thereby maintaining the final overlapping uORF with the main ORF for *fluc* (Promega). HEK293 cells were plated in 24-well plates at a density of 25,000 cells/well and incubated overnight. *Fluc* reporter plasmids and pRL-TK renilla luciferase (rluc) transfection control plasmid (Promega) were co-transfected into these cells using jetPRIME® transfection reagent (Illkirch-Graffenstaden, France). Cells were incubated overnight in the presence or absence of 1 μM thapsigargin (TG) (Sigma). Cells were then lysed and the Fluc/Rluc ratio was determined using the Promega Dual-Luciferase Reporter kit (Madison, USA).

### Immunodetection

Cultured satellite cells were fixed in 4% paraformaldehyde (PFA), permeabilized with 0.2% Triton, 50 mM NH_4_Cl and blocked in 5% horse serum (HS). Single EDL myofibers were fixed with 4% PFA, permeabilized with 0.1% Triton in PBS and blocked in 5% HS with 0.1% Triton in PBS. TA muscles were fixed for 2 hr in 0.5% PFA at 4°C and equilibrated overnight in 20% sucrose at 4°C. Tissues were mounted in Frozen Section Compound (VWR) and flash frozen in a liquid nitrogen-cooled isopentane bath. Transverse sections (10μm) were permeabilized with 0.1% Triton, 0.1M Glycine in PBS, and blocked in M.O.M. reagent (Vector Labs). For immunoblotting, cell lysates were obtained in RIPA buffer (ThermoFisher Scientific) supplemented with Complete protease inhibitor cocktail (Roche) and phosphatase inhibitor cocktail (Sigma).

Primary antibodies were against PAX7 (DSHB), MYOD (SantaCruz, sc-304 and sc-377460), TACC3 (Abcam 134154), Ki67 (BD Biosciences B56), LAMININ (Sigma L9393), MYOGENIN (Abcam 124800) and β-ACTIN (Sigma A5441). Alexa Fluor-488 and Alexa Fluor-594 conjugated secondary anti-mouse IgG1, anti-mouse IgG2B or anti-rabbit antibodies (Life Technologies) were used for immunofluorescence. Horseradish peroxidase (HRP) conjugated anti-mouse or anti-rabbit secondary antibodies (Jackson Immunoresearch) were used with the ECL Prime Western Blotting Detection reagents (GE Healthcare). Densitometry of immunoblots was performed with ImageJ.

### RNA Analysis

RNA was isolated from cells with TRIzol reagent (Life Technologies) and treated with DNase (Roche) prior to reverse transcription with iScript reverse transcription supermix (BioRad). RT-PCR primers were *Pax7* forward 5’-AGGCCTTCGAGAGGACCCAC-3’ reverse 5’-CTGAACCAGACCTGGACGCG-3’, *Myogenin* forward 5’-CAACCAGGAGGAGCGCGATCTCCG-3’ and reverse 5’-AGGCGCTGTGGGAGTTGCATTCACT-3’, *Tacc3* forward 5’-GAGCTTCAGAGACCCATCAGA -3’ reverse 5’-AGTTGGAGAGATGGGACGAG-3’ and *Actb* forwad 5’-AAACATCCCCCAAAGTTCTAC-3’ and reverse 5’-GAGGGACTTCCTGTAACCACT-3’. Levels of mRNA were measured using SYBR Green on a 7500 Fast Real Time PCR System (Applied Biosystems).

### RNA Sequencing, Mass Spectrometry and Bioinformatic Analysis

For both RNA and protein analysis, satellite cells were isolated from *Pax3^GFP/+^* adult mice and seeded at 7500 cells per 35mm plate and subsequently cultured in 10 μM sal003 (n=4) or DMSO (n=4) for 4 days. For RNA analysis, total RNA was isolated from satellite cell cultures using the RNeasy® Micro Kit (Qiagen). The RiboGone™-mammalian (Takara Bio) kit was subsequently used to eliminate ribosomal and mitochondrial RNA. cDNA libraries for RNA-sequencing were then prepared using the SMARTer® Stranded RNA-Seq (Takara Bio) kit. With all kits, the manufacturer’s protocols were followed. The resulting cDNA libraries were sequenced with the Illumina MiSeq system (Genome Quebec).

For protein analysis, the TMTsixplex™ Isobaric Tag Kit (Thermo Fisher Scientific) was used to isolate and label peptides from satellite cells cultured in 10 μM sal003 (n=12 plates), or DMSO (n=12 plates), per the manufacturer’s protocol. After labelling peptides with unique isobaric tags, the peptide solutions were pooled and analysed via high pH reversed phase fractionation and LC-MS/MS. All data was acquired with Thermo Orbitrap Fusion™ Tribrid™ 2.1 software (Thermo Fisher Scientific), analyzed using Proteome Discoverer 1.4 (Thermo Fisher Scientific) and MASCOT v2.4 software (Matrix Science). Raw data files were searched against a Uniprot Mouse database. Gene lists were examined using Enrichr(18) for ENCODE and ChEA consensus transcription factor binding sites, ENCODE histone modifications, mouse tissue expression, gene ontology (GO) Biological Process and GO Cellular Component. Venny2.1.0 was used to generate a list of genes upregulated at the protein level (log2>1), without a corresponding increase in mRNA expression (log2<1).

### Computational analysis

Trimmomatic v0.32(35) was used to trim sequencing reads, including adaptors and other Illumina-specific sequences, the first four bases from the start of each read and low quality bases identified using a 4bp sliding window where quality fell below 30 (phred33 < 30). Finally, reads shorter than 30 base pairs were removed. Cleaned reads were aligned to the mouse reference genome build mm10 using STAR v2.3.0e(36) with default settings. Reads mapping to more than 10 locations in the genome (MAPQ < 1) were discarded. Gene expression levels were estimated by quantifying uniquely mapped reads to exonic regions (the maximal genomic locus of each gene and its known isoforms) using featureCounts (v1.4.4)(37) and the Ensembl gene annotation set. Normalization (mean of ratios) and variance-stabilized transformation of the data were performed using DESeq2 (v1.14.1)(38). Multiple control metrics were obtained using FASTQC (v0.11.2), samtools (v0.1.20)(39), BEDtools (v2.17.0)(40) and custom scripts. For visualization, normalized Bigwig tracks were generated using BEDtools and UCSC tools. Integrative Genomics Viewer(41) was used for data visualization. PCA was done using the 1000 most variant genes.

### Statistical analysis

Graphs are presented as mean ± SEM, as indicated in Figure legends. Unless otherwise indicated, three independent replicates of each experiment were performed. Significance was calculated by unpaired Student’s t tests with two-tailed p values: *p < 0.05, **p < 0.01, ***p < 0.001.

## Acknowledgements

We thank C. Young for assistance with flow cytometry, C. Borchers and D. Smith for assistance with mass spectrometry. R. Yao generously provided *Tacc3^fl/fl^* mice. C.C. and coworkers are funded by the Canadian Institute for Health Research (CIHR 399258), the Stem Cell Network, the Fonds de Recherche du Québec en Santé (FRQS) and the Strauss Foundation. R.F. is funded by JSPS Overseas Research Fellowship, the Uehara Memorial Foundation, and the Mochida Memorial Foundation for Medical and Pharmaceutical Research. C.L.K. is supported by the CIHR (156086) and the FRQS. Data analyses were enabled by computer and storage resources provided by Compute Canada and Calcul Québec.

---

**Supplementary Fig. 1.**
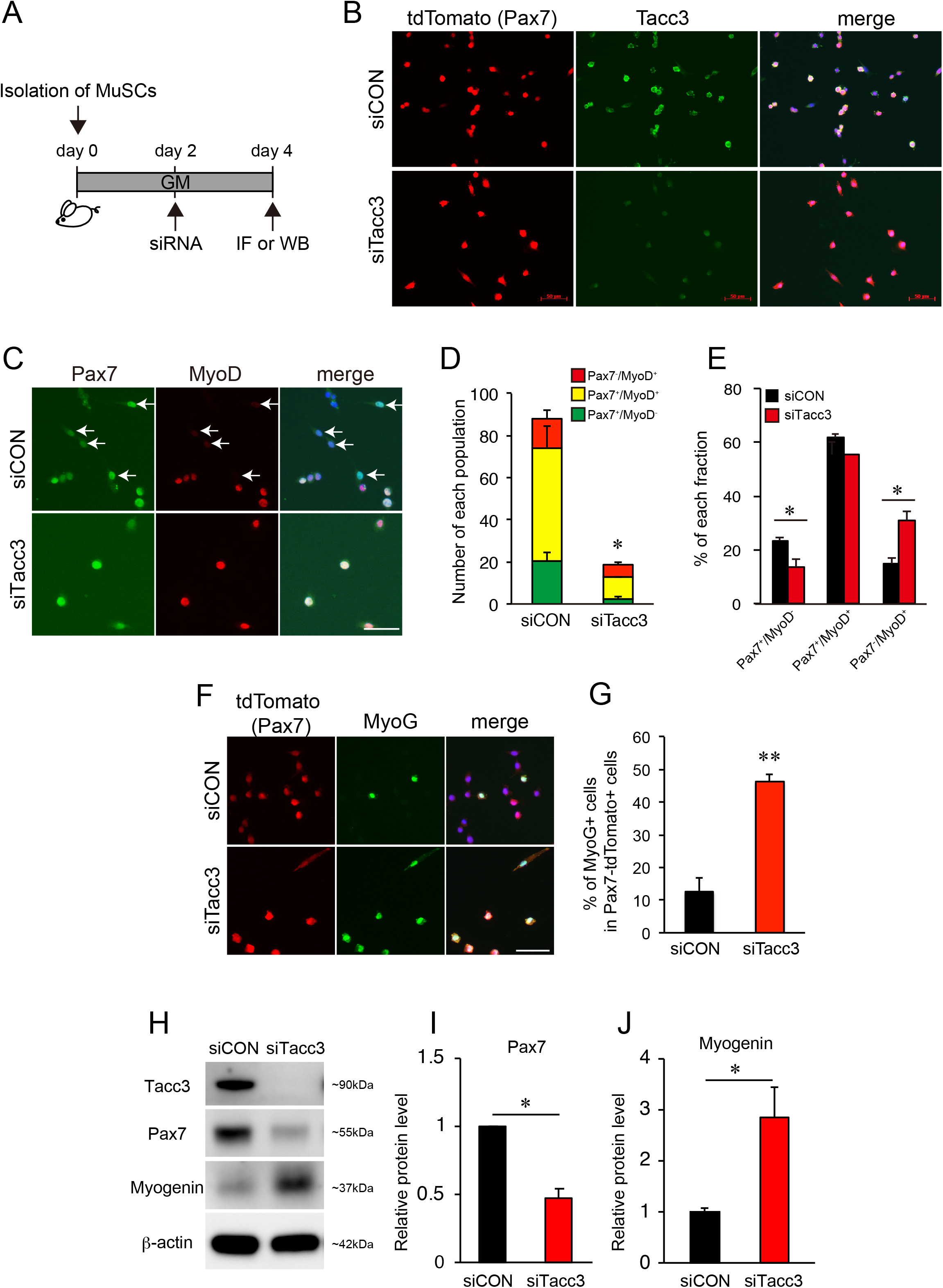
*Tacc3* knockdown limits satellite cell expansion and self-renewal *ex vivo*. (*A*) Satellite cells were isolated from tmx treated Pax7CreERT2/+; R26tdtomato/+ mice and transfected with siRNAs against *Tacc3*. (*B*) Tdtomato fluorescence (PAX7, red) combined with immunofluorescence analysis with antibodies against TACC3 (green) after 4-day culture of satellite cells transfected (day 2) with siRNA against *Tacc3* (siTacc3) and control siRNAs (siCON). (*C*) Immunofluorescence analysis with antibodies against PAX7 (green) and MYOD (red) after 4-day culture of satellite cells transfected (day 2) with siRNA against *Tacc3* (siTacc3) and control siRNAs (siCON). (*D-E*) Quantification of numbers of PAX7(+), MYOD(−) satellite cells undergoing self-renewal (reserve cell population), activated PAX7(+), MYOD(+) satellite cells and differentiating PAX7(−), MYOD(+) satellite cells in (C). (*F*) Tdtomato fluorescence (PAX7, red) combined with immunofluorescence with antibodies against MYOGENIN (green) after 4-day culture of satellite cells transfected (day 2) with siRNA against Tacc3 (siTacc3) and control siRNAs (siCON). (*G*) Quantification of numbers of PAX7(+) nuclei and MYOGENIN(+) nuclei in (F). (*H*) Immunoblotting of lysates with antibodies against PAX7, MYOGENIN, TACC3 and β-actin, after 4-day culture of satellite cells transfected (day 2) with siRNA against Tacc3 (siTacc3) and control siRNAs (siCON). Quantification of (*I*) PAX7 and (*J*) MYOGENIN levels in (H) by densitometry. Levels are normalized to β-actin. Scale bars represent 50μm (B, C and F). Values indicate mean (n = 3) ± SEM * p < 0.05, ** p < 0.01.

**Supplementary Fig. 2.**
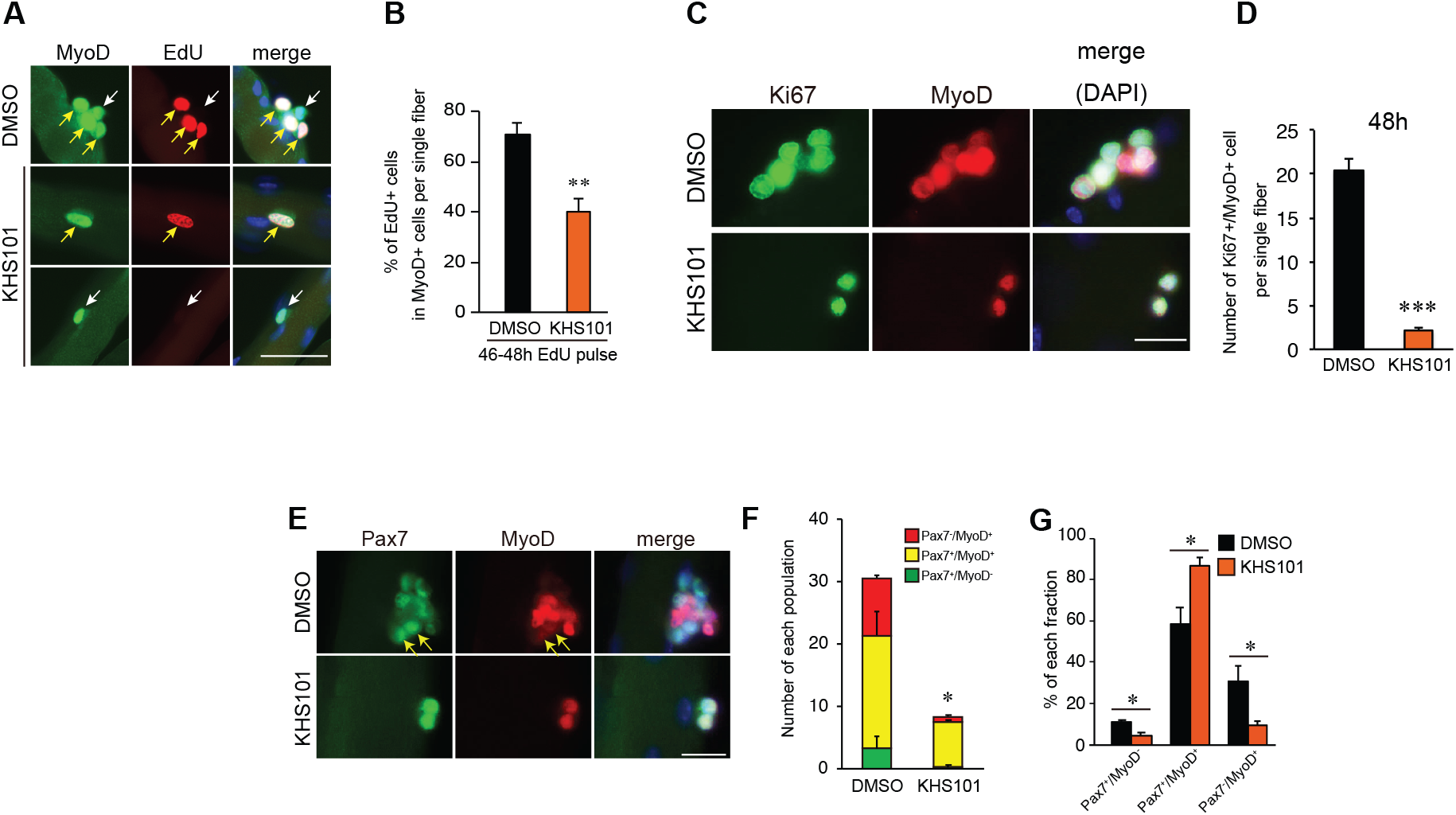
Pharmacological inhibition of TACC3 limits satellite cell expansion and self-renewal *ex vivo*. (*A*) Immunolabeling with antibodies against MYOD (green) and EdU incorporation (red) on single EDL myofibers cultured in the presence of TACC3 inhibitor KHS101 for 48 hours. (*B*) Percent of EdU(+), MYOD(+) satellite cells in (A). (*C*) Immunolabeling with antibodies against MYOD (green) and Ki67 (red) on single EDL myofibres cultured in the presence of TACC3 inhibitor KHS101 for 48 hours. (*D*) Percent of Ki67(+), MYOD(+) satellite cells in (C). (*E*) Immunolabeling with antibodies against PAX7 (green) and MYOD (red) on single EDL myofibres cultured in the presence of TACC3 inhibitor KHS101 for 48 hours. (*F-G*) Quantification of numbers of PAX7(+), MYOD(−) satellite cells undergoing self-renewal (reserve cell population), activated PAX7(+), MYOD(+) satellite cells and differentiating PAX7(−), MYOD(+) satellite cells in (E). Scale bars represent 50μm (A) or 20μm (C, E). Values indicate mean (n = 3) ± SEM * p < 0.05, ** p < 0.01, *** p < 0.001.

**Supplementary Data 1.** Transcriptome analysis of satellite cells cultured in the presence of 10μM sal003.

**Supplementary Data 2.** Identification and quantitation of proteins in satellite cells cultured in the presence of 10μM sal003 using tandem mass spectrometry.

**Supplementary Data 3. Satellite cells upregulate protein levels independent of mRNA when cultured in sal003.** 140 genes that are upregulated at the level of protein (mass spectrometry, log2>1), without a corresponding increase reported by mRNA (RNA-seq, log 2<1) are organized with respect to selected biological processes.

